# Rhinovirus 2A is the predominant protease responsible for instigating the early block to gene expression encountered in infected cells

**DOI:** 10.1101/591032

**Authors:** David Smart, Irene Filippi, Benjamin Smalley, Donna Davies, Christopher J. McCormick

## Abstract

Human rhinoviruses express 2 cysteine proteases, 2A and 3C, that are responsible for viral polyprotein processing. Both proteases also suppress host gene expression by inhibiting mRNA transcription, nuclear export and cap-dependent translation. However, the relative contribution that each makes in achieving this goal remains unclear. In this study we have compared both the combined and individual ability of the 2 proteases to shut down *in cellulo* gene expression using a novel dynamic reporter system. Our findings show that 2A inhibits host gene expression much more rapidly than 3C. By comparing the activities of a representative set of proteases from the three different Human Rhinovirus (HRV) species, we also find variation in the speed at which host gene expression is suppressed. Our work highlights the key role that 2A plays in early suppression of the infected host cell response and shows that this can be influenced by natural variation in the activity of this enzyme.

## Introduction

Human rhinoviruses typically infect the upper respiratory tract, and are the most common etiological agent responsible for the common cold. Infections are acute, and usually cause only minor symptomatic issues. However, for individuals who have asthma, and the in elderly, infections are more problematic, with these groups experiencing notable levels of morbidity and even mortality[1]. As a result, HRV infection places a significant health burden on society that in North America alone has an estimated economic impact of around $40 billion[2]. For this reason, there is a need to understand in detail the drivers of HRV–induced pathology to better target intervention strategies.

Rhinoviruses belong to the *enterovirus* genus, and are themselves subdivided into 3 genetically distinct species (HRV-A, B and C)[3]. All enteroviruses express two separate proteases, 2A and 3C, which are used to usurp infected host cell functions. Each protease targets different host cell substrates, as well as being responsible for processing different boundaries within the viral polyprotein. In part due to the requirement placed on it to process the majority of polyprotein boundaries, 3C has a well-defined substrate recognition sequence but one which still allows it to target proteins such as poly-A binding protein [4, 5], Oct-1[6], TATA-binding protein [7, 8], CstF-64[9] and nucleoporin 153 (Nup153)[10]. In contrast, 2A needs only to cleave itself away from the viral polyprotein at its amino terminal boundary. Possibly as a consequence of this, the protease has a less well-defined recognition motif, with cleavage of some host substrates depending not just on active site recognition but also exosite interactions as well[11, 12]. Host proteins targeted by 2A include eIF4GI[13], elF4GII[14], Gemin3[15] and several nucleoporins (Nup62, 98 and 153)[16, 17]. Overall, the cleavage of host proteins by 2A and 3C inhibits mRNA transcription, processing, export and translation, and represents a key strategy used by the virus to prevent the cell mounting a successful antiviral response. However, the relative contribution that cleavage of each host substrate makes to blocking the host cell’s potential for new gene expression remains unclear. Certainly, 2A-dependent cleavage of both forms of eIF4G correlates with the early shutdown of global cap-dependent mRNA translation in the infected cell[14, 18]. However, inhibition of gene expression from recently transcribed mRNAs occurs at an even earlier time point and correlates with a reduction in mRNA transport from the nucleus, pointing to additional involvement of cleavage of other proteins such as Nups[19, 20]. Other studies have shown that expression of a genetically engineered TATA-binding protein resistant to 3C cleavage suppresses poliovirus replication[8], consistent with the notion that the reduction in transcription driven by this protease may play a contributory role in early suppression of potential antiviral responses. Indeed, enteroviruses rely on active transport to direct 3C into the nucleus at early time points, when it is predominantly present as a 3CD precursor[21, 22], suggesting early protease activity in this organelle is important for infection.

There is variability both within and between HRV species in the ability of 2A to cleave its substrates. Early studies found that 2A from HRV-2A cleaves eIF4GI and eIF4GII at approximately similar rates[23] whereas the same protease from HRV-B14 displays a substrate preference for eIF4GI[18]. More recently it has been confirmed that there is variation between all 3 HRV species regards the cleavage of both eIF4GI and Nup [17]. Furthermore, this variation in Nup cleavage potentially correlates with differences in the speed at which 2A disrupts nuclear import and export of fluorescent reporter proteins, and by inference mRNA export[24]. It is therefore possible that there is an inherent difference between different HRV proteases in the rate at which they block new gene expression. However, measuring whether differences exist is challenging. Firstly, the multifaceted nature by which the proteases inhibit gene expression means that analysis has to be done using cells rather than *in vitro* experimentation. Complications also arise from the use of infectious virus to drive protease expression, in part because of differences in the rate of genome replication between different strains, which in turn dictates rates of protease expression. Expression of proteases from DNA or RNA constructs transfected into cells offers an alternative solution to this latter problem. However, such an approach requires that changes to new gene expression be detected before the proteases introduce unwanted experimental variability through significantly altering their own expression.

In this study, we have used a dual promoter mammalian expression construct co-expressing HRV proteases alongside a luciferase-reporter gene to examine potential differences in the rate at which 2A and 3C switch off new gene expression. Importantly, both the reporter protein and its mRNA have been engineered for rapid turn-over, thus generating a system that is highly responsive to early changes imposed by the proteases. We find that there are detectable differences when comparing between representative proteases from the three different HRV species. Furthermore, the early rapid shut-down of gene expression we observe appears to be exclusively due to 2A with 3C playing little or no role.

## Methods

### Cell lines and reagents

293T cells were maintained at 37°C in a 5% CO_2_ incubator with DMEM (Invitrogen, Fisher Scientific UK Ltd, Loughborough, UK) supplemented with 10% foetal bovine serum, 100 U/ml penicillin, 100 μg/ml streptomycin. Experimental treatment of cells included incubation in medium supplemented with 200 μg/ml cycloheximide (Sigma, Gillingham, UK) and/or 5 μg/ml actinomycin D (Cayman Chemical Cambridge Bioscience Ltd, Cambridge, UK).

### DNA constructs

Generation of some constructs relied on gene synthesis (Invitrogen, Fisher Scientific UK Ltd, Loughborough, UK). In instances where this was the case, sequences ordered were codon optimized for mammalian expression and then manually adjusted to minimize the existence of unwanted splice donor and splice acceptor sites using the online programmes HSF3[25] and the NetGene2 server[26]. The initial plasmid expressing GFP, VSV-tagged 2A and HA-tagged 3C from HRV16 was generated by cloning a single synthetic DNA fragment into an in-house dual promoter plasmid via *EcoRI* and *SalI* restriction sites to generate pCIPEP-A16^TAG^(+/+) (see Fig. S1 for sequence). Synthesized DNAs encoding for HRV-B4 and HRV-C2 proteases (Table S1) were exchanged with their respective counterparts in pCIPEP-A16^TAG^(+/+) using ClaI and *SbfI* (2A) and *BsiWI* and SalI (3C) restriction sites. Inactivation of the proteases involved a two-step PCR approach, which introduced a Cys>Ala mutation in the active site of both 2A and 3C. A similar PCR-based strategy was used to generate constructs expressing tag-free proteases and a 3C protease with a c-Myc nuclear localization signal (PAAKRVKLD) fused directly to its C-terminus. Primer sequences are available on request.

The synthetic DNA encoding for the Thioredoxin-Nanoluciferase(Nluc)PEST fusion protein linked by a peptide derived from translating the hepatitis delta virus (HdV) ribozyme sequence (HdV^WT^Nluc; Fig. S2) was initially cloned into pCDNA3.1 via *XbaI* and *BamHI* restriction sites. Introduction of synonymous mutations to block the HdV ribozyme activity was achieved by a two-step PCR approach with the final product (HdV^KO^Nluc) cloned back into pCDNA3.1. Subsequently, both Nluc-containing coding regions were transferred from pCDNA3.1 to the pCIPEP vectors using *BamHI* and *XbaI* restriction sites. The other luciferase encoding vector, pR.EMCV.F, used to assess cap-dependent vs. cap-independent translation, has been described previously[27]. All constructs created for the purpose of this project are freely available on request.

### Transfections

For transfection of adherent cell monolayers, cells were seeded at a density of 4 × 10^4^ cm^2^. The next day they were transfected over 24 hours with a DNA-Fugene HD mixture at a ratio of 1 μg DNA to 3 μl Fugene (Promega, Southampton, UK) according to the manufacturer’s instructions (Western analysis) or using a DNA:polyethylenimine ratio of 3:1 (Northern blot and Luciferase assays)[28].

For transfection of cell suspensions by electroporation, cells were detached with trypsin/EDTA, washed × 2 using ice-cold serum-free RPMI (Invitrogen, Fisher Scientific UK Ltd, Loughborough, UK) and resuspended in RPMI at a final viable cell density of 1.5 × 10^7^ ml^-1^. Unless otherwise stated, 400 μl of this cell suspension was mixed with 5 μg plasmid DNA, transferred to a pre-chilled 0.4 cm cuvette and electroporated at 270 V, 950 uF using a BioRad GenePulser II. Cells were subsequently flushed out into 2.8 ml ice-cold growth medium and the suspension left on ice until all transfections within the experiment had been completed. Two hundred and fifty microliters of the cell suspensions were then used to seed wells of a 12-well plate, already containing 1 ml fresh growth medium pre-equilibrated to 37°C. Stochastic variation between transfection efficiencies were often observed within a single experiment, based on luciferase readouts at 1 hour. For this reason, each repeat was carried out with new batches of plasmid DNAs, the order in which these DNA constructs were transfected was randomized, and a statistical assessment of transfection efficiencies across all experiments (based on unadjusted 1 hour reads) undertaken to ensure no obvious transfection-based bias remained.

### Luciferase assays

Cells were detached by vigorous pipetting, pelleted by centrifugation at 500g for 1 min at 4°C, lysed in 100 μl Passive Lysis Buffer (Promega, Southampton, UK) and stored at −70°C. Upon thawing of lysates, luciferase activity was determined using either the Dual Luciferase Assay System (Promega) or Nano-Glo Luciferase Assay System (Promega) according to manufacturer’s recommendations.

### Western blot

Cells were detached, washed in phosphate buffered saline, and lysed in RIPA buffer (50mM Tris-HCl pH8.0, 150mM NaCl, 1%[w/v] NP-40, 0.5% [w/v] sodium deoxycholate, 0.1% [w/v] sodium dodecyl sulphate) supplemented with 2 × cOmplete™ protease inhibitor (Sigma, Gillingham, UK). After centrifugation at 14,000g for 1 minute to pellet cell nuclei, supernatant removed and collected, and assessed by BCA assay (Pierce, Fisher Scientific UK Ltd, Loughborough, UK) to determine protein concentrations. Samples (10 μg/well) were run on an sodium dodecyl sulphate-polyacrylamide gel and transferred to polyvinylidene fluoride membranes (Amersham) which were then probed with primary antibodies against AU-rich element RNA-binding protein 1 (AUF1) (D604F; Cell Signalling, London, UK), hemagglutinin (HA) tag (16B12; Biolegend UK Ltd, London, UK) glyceraldehyde 3-phosphate dehydrogenase (GAPDH) (mAb374; Chemicon, Sigma, Gillingham, UK), eIF4GI (D6A6; Cell Signalling, London, UK), green fluorescent protein (GFP) (Serotec, Bio-Rad, Watford, UK) and vesicular stomatitis virus (VSV) tag (Biolegend UK Ltd, London, UK) followed by the appropriate secondary peroxidase-conjugated antibody (Sigma, Gillingham, UK). Bound antibody was detected using Clarity ECL Western Blotting Substrate (Bio-Rad, Watford, UK) with the image digitally captured using an Amersham Imager 600 (GE Healthcare Life Sciences, Little Chalfont, UK).

### Northern blot

RNA was recovered from transfected cells using TriFAST Reagent (Peqlab, Lutterworth, UK) according to the manufacturer’s instructions. Purified RNA was run on a 0.8% MOPS-formaldehyde agarose gel, stained using SybrGold (Invitrogen, Fisher Scientific UK Ltd, Loughborough, UK) to confirm rRNA integrity and subsequently transferred to charged nylon membranes. After UV-crosslinking, membranes were pre-blocked by a 30 minute incubation in Ultrahyb (Invitrogen, Fisher Scientific UK Ltd, Loughborough, UK) at 42°C, before a biotinylated DNA probe, generated by PlatinumBrightBIO (Kreatech, Leica Microsystems (UK) Ltd, Milton Keynes, UK), was added. Probes used included a cDNA fragment from the protease encoding pCIPEP transcript (nucleotides 122 to 1580 of the predicted mRNA transcript encompassing the hepatitis C virus (HCV) internal ribosome entry site (IRES), the GFP coding region and the encephalomyocarditis virus (EMCV) IRES), as well as the *XbaI-BamHI* TNP^HdV+^ cassette. After overnight incubation and washing of the membrane at 42°C, firstly in 2xSSPE + 0.1% SDS and subsequently in 0. 1XSSPE 0.1% SDS, bound probe was detected using the BrightStar Northern Blot detection kit (Invitrogen, Fisher Scientific UK Ltd, Loughborough, UK). A GAPDH probe was used on parallel blots (Nluc assay experiments) or reprobed blots (HRV protease transcript experiments) to confirm loading and RNA integrity; the decision to use parallel blots being based on probe signal overlap coupled with an inability to strip membranes when using BrightStar detection. Images were captured on film.

### Immunofluorescent Imaging

Cells seeded on glass coverslips were transfected using Fugene HD and subsequently fixed with ice-cold 4% paraformaldehyde in phosphate buffered saline. Fixed cells were blocked and permeabilized using 10% foetal calf serum, 1% saponin in phosphate buffered saline for 2 hours, stained with a 1:1000 anti-HA mAb and 1:100 anti-mouse tetramethylrhodamine (TRITC) secondary antibody (Sigma), mounted using a 4,6-diamidino-2-phenylindole (DAPI) containing mountant (Vectashield, Vector Laboratories Ltd, Peterborough, UK) and visualized using a Leitz DMRB fluorescence microscope with a 100x oil immersion lens.

### Statistical analysis

Pairwise comparisons were made using a paired Students t-test. Analysis of three or more experimental groups was undertaken using a repeated measures ANOVA applying Greenhouse-Geisser correction to identify within-subject effect differences and adjusting for multiple comparisons using Bonferroni when undertaking subsequent pairwise comparisons.

## Results

### Producing a co-expression system for 2A and 3C

A series of PolII-based constructs was generated that produced a tricistronic mRNA where translation of the first cistron was under the control of an HCV IRES and translation of the latter two cistrons were under the control of EMCV IRESes. GFP was placed within the first cistron while the second and third cistrons encoded an N-terminal VSV-tagged 2A and N-terminal HA-tagged 3C protease respectively, both of which were expressed as ubiquitin fusion products (Fig. 1a). Inclusion of the ubiquitin coding regions served two purposes. They distanced the 2A and 3C coding regions from their respective start codons, so that any change in protease sequence would have minimal influence on translation initiation[29]. They also offered the potential for expression of 2A and 3C lacking N-terminal extensions if required.

**Figure 1.**
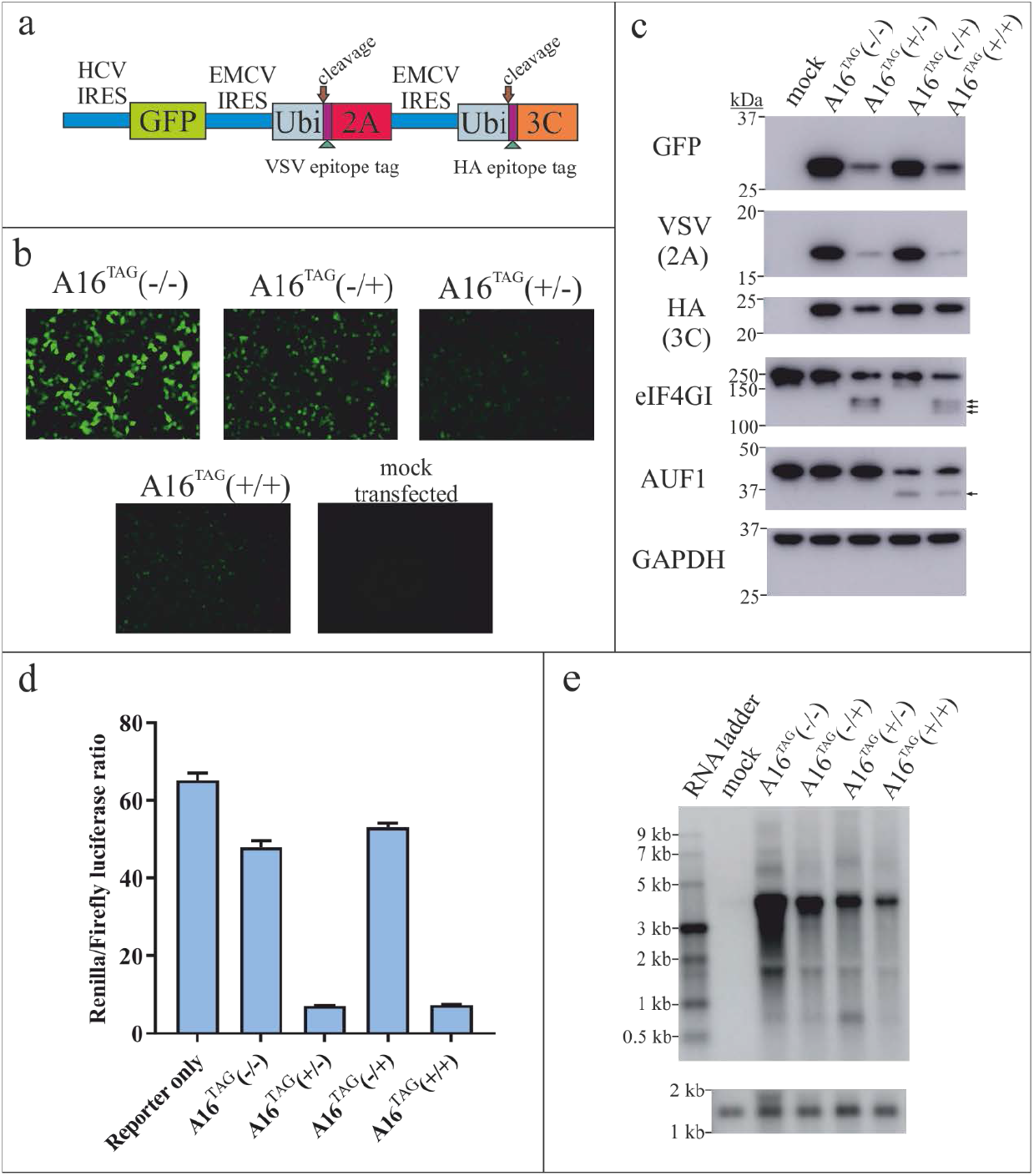
Combined expression of GFP and epitope-tagged HRV A16 proteases from a series of tricistronic expression constructs. (a) A schematic outlining expression construct design. (b) Visualization of GFP in cells transfected with the different A16 epitope tagged expression constructs. (c) Western blot of transfected cells. Arrows indicate the position of protease cleavage products. (d) Dual luciferase ratios obtained from cells co-transfected with the protease expression constructs and a dual luciferase assay reporter (pR.EMCV.F) encoding Renilla and Firefly luciferases under the control of 5’cap and EMCV IRES respectively (data represents the mean ± S.E.M from triplicate readings). (e) Transcript production assessed by Northern blot (e) using a probe complementary to the protease-encoding transcript (upper panel) or a GAPDH probe (lower panel).

Initially four constructs were generated that expressed epitope-tagged HRV-A16 2A and 3C proteases; the first was designed to express two inactive proteases (A16^TAG^(−/−)), the second two active proteases (A16^TAG^(+/+)), the third an inactive 2A and active 3C (A16^TAG^(−/+)) and the fourth an active 2A and inactive 3C (A16^TAG^(+/−)). Visualization of 293T cells transfected with these constructs showed that all four expressed GFP to detectable levels (Fig 1b). Western blot analysis of transfected cell lysates (Fig. 1c) confirmed expression of GFP, and verified that there was production of HA and VSV tagged proteins that were of the size expected for 2A and 3C respectively. As would be expected with the introduction of functional HRV proteases into cells, all constructs encoding for one or more active protease had reduced expression of their proteins, although the impact that each protease had was noticeably different. Compared with A16^TAG^(−/−), A16^TAG^(−/+) transfected cells exhibited only a very modest reduction in expression of GFP, 2A and 3C. In contrast, despite anticipating that 2A would have a composite effect of enhancing IRES-dependent translation from the tricistronic transcript while reducing transcript production, A16^TAG^(+/−) cells exhibited a more marked reduction in expression of all three antigens. This latter profile was similar to that seen for A16^TAG^(+/+).

To verify the activity of the proteases, the same Western blots were probed with antibodies against eIF4GI and AUF1, substrates for 2A and 3C respectively[13, 30]. Cleavage of eIF4GI was only seen in A16^TAG^(+/−) and (+/+) lysates whereas a reduction in AUF1 expression was only observed in A16^TAG^(−/+) and (+/+) lysates (Fig. 1c). A faint AUF1 3C cleavage product was also occasionally observed, likely representing a labile cleavage product. Combined cleavage of eIF4GI and II by 2A should lead to a shutdown of cap-dependent translation while enhancing IRES-dependent translation[18]. To examine this, the protease expressing constructs were co-transfected into cells with a bicistronic reporter plasmid that expressed renilla and firefly luciferase through a cap and EMCV-IRES dependent mechanism respectively (Fig. 1d). A decrease in the ratio of renilla:firefly was only seen in A16^TAG^(+/−) and (+/+) transfected cells, consistent with active 2A targeting both forms of eIF4G and shutting down cap-dependent translation.

To verify transcript production was occurring as expected, RNA from cells transfected with each of the A16^TAG^ constructs was analysed by Northern blot (Fig. 1e). One major transcript of the expected size was seen, consistent with translation of GFP, 2A and 3C being from a single RNA. Levels of this RNA also varied between the experimental groups in a manner suggesting that both proteases were able to suppress transcript levels.

### Extending the expression system to include HRV-B and HRV-C proteases

To allow comparisons between different HRV species proteases, the A16 protease coding regions were exchanged with those from HRV-B4 and HRV-C2, generating B4^TAG^(+/+), B4^TAG^(−/−), C2^TAG^(+/+) and C2^TAG^(−/−). Northern blot analysis of 293T cells transfected with these constructs alongside the original A16^TAG^ constructs verified that a single major transcript of the expected size was produced from each (Fig. 2a). Furthermore, the abundance of this transcript was similar when comparing between all (−/−) constructs and when comparing between all (+/+) constructs, although consistent with earlier Northern blot analysis, expression of the active proteases did reduce transcript abundance.

**Figure 2.**
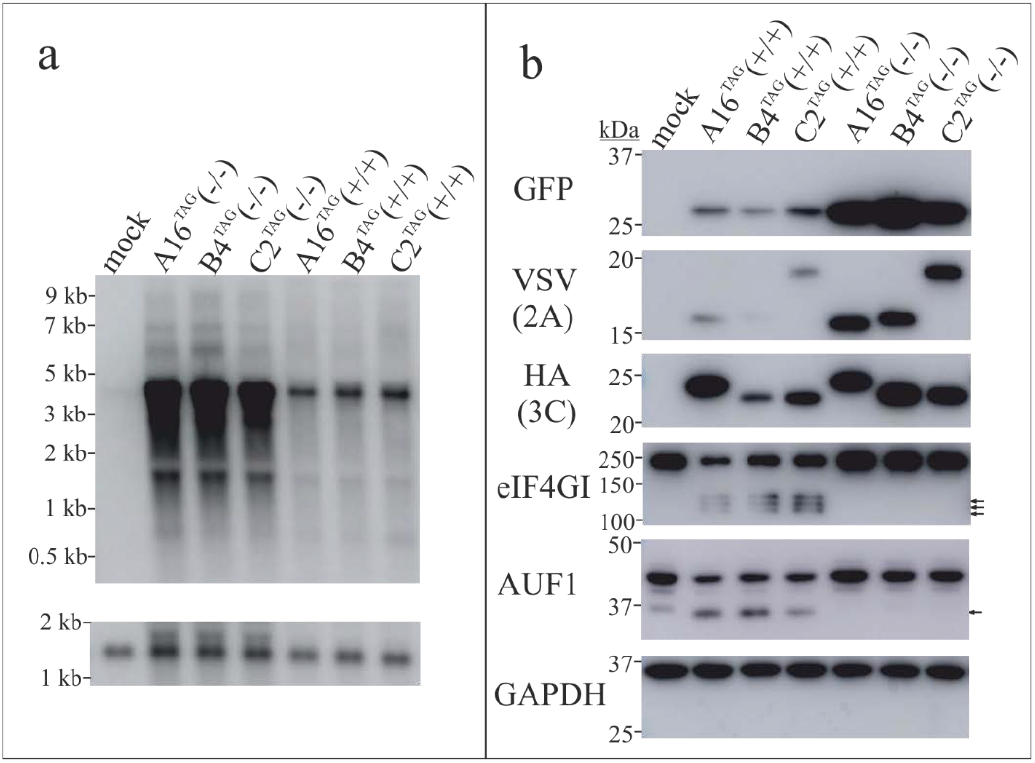
Expression of epitope tagged 2A and 3C for all three HRV species. (a) A Northern blot of cells transfected with different tricistronic protease expression constructs and hybridized to a probe complementary to the protease-encoding transcript (upper panel) or GAPDH (lower panel). (b) Western blot analysis of cells transfected with the same constructs. Arrows indicate the position of protease cleavage products.

Western analysis confirmed that all constructs expressed GFP, VSV-tagged 2A and HA-tagged 3C (Fig. 2b). Importantly, cleavage of AUF1 and eIF4G1 was seen in all (+/+) lanes, verifying that each of the three different 2A and 3C proteases were being expressed in an active form. While the levels of GFP, 2A and 3C were far higher in all (−/−) lanes compared to all (+/+) lanes, differences were also observed when comparing within the (+/+) or (−/−) groups. One of the more subtle differences was that seen for GFP expression, with this protein being slightly lower and slightly higher in the B4(+/+) and B4(−/−) group respectively compared to the other protease active and inactive constructs. Expression of 3C was similar when comparing between (−/−) groups but showed reduced expression in the B4(+/+) experimental group compared to the other (+/+) groups. Interestingly, expression levels of 2A were more varied compared to GFP and 3C, and were less consistent between experiments, particularly when comparing between the different (+/+) groups (Fig. S3). Prior indirect evidence suggest that 2A stability may vary to some degree between isolates[24] and based on our data (Fig. S3) HRV-C2 2A might also have been subject to partial proteolytic cleavage in some of the experiments. Differing 2A half-lives coupled to the increasingly complex and less predictable environment which this protein and 3C find themselves in as more and more cellular functions are disrupted by protease activity may well be driving the patterns of variation seen. Nonetheless, based on the (−/−) data, we conclude that before such forces manifest themselves, protease expression from the different constructs occurs in a manner that is broadly consistent and reproducible.

### Development of a reporter system for dynamic monitoring of changes to gene expression

A highly dynamic reporter system exhibiting rapid turnover of both mRNA and protein should allow shut down of new gene expression by the HRV proteases to be detected before this event has a pronounced impact on HRV protease expression itself. However, the intrinsic mechanisms adopted by the cell to limit mRNA half-life, such as the positioning of AU rich regions within mRNA 3’ untranslated regions and nonsense-mediated decay, are themselves subject to regulation and often manipulated by viruses[30, 31]. It was therefore desirable to have a system where mRNA was destabilised in a manner that was independent of normal host regulatory pathways. To achieve this a destabilized Nluc coding region was linked to an upstream thioredoxin coding region via a Hepatitis Delta Virus (HdV) ribozyme sequence, thus generating a single continuous ORF that should be subject to internal cleavage (Fig 3a). An intron was placed within the HdV ribozyme sequence to minimize ribozyme cleavage during mRNA transcription as well as ensure that RNA splicing had to occur for luciferase activity to be observed.

**Figure 3.**
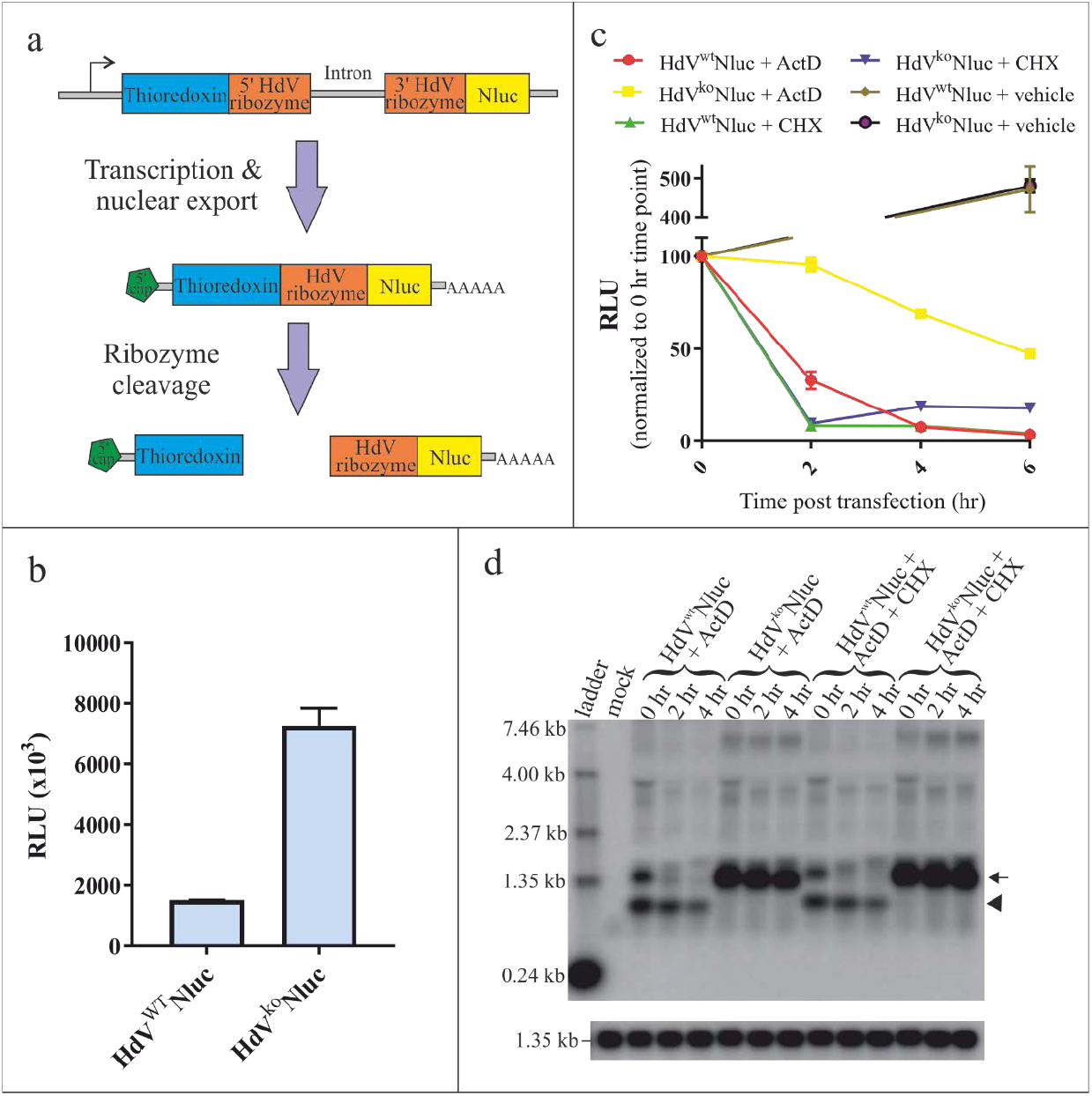
Validating the use of a dynamic luciferase reporter expression cassette. (a) A schematic of the HdV^WT^Nluc reporter cassette and its expected processing, with ribozyme cleavage of the mRNA leading to translational termination of Nluc. (b) Relative luciferase activity (measured in relative light units (RLU)) from cells transfected with a plasmid expressing either a functional (HdV^WT^Nluc) or inactive (HdV^ko^Nluc) ribozyme reporter cassette (values represent the mean ± s.d. from 2 separate experiments). (c) Relative changes in luciferase values after treatment of transiently transfected cells with either CHX or ActD (n=2; values represent the mean ± s.d.). (d) Northern blot of RNA from transiently transfected cells treated with ActD, or ActD + CHX. The blot in the upper panel has been hybridized to a probe derived from the HdV^WT^Nluc ORF, while that in the lower panel has been hybridized to a GAPDH control probe. The arrow and arrow head indicate the predicted size of the intact reporter mRNA its 3’ cleavage product respectively.

To test reporter gene expression from this cassette, a plasmid carrying this active ribozyme-embedded ORF (HdV^wt^-Nluc) or a comparable control ORF encoding for a defective ribozyme (HdV^ko^-Nluc) were transfected into cells. Consistent with the HdV ribozyme destabilizing the RNA it was embedded in, HdV^wt^-Nluc transfected cells exhibited 5-fold lower luciferase activity compared to HdV^ko^-Nluc transfected cells (Fig. 3b). Cells were also transfected with the same constructs and then treated with either cycloheximide (CHX) or actinomycin D (ActD). Monitoring of luciferase activity (Fig. 3c) showed that the CHX-imposed block to translation resulted in an early rapid drop in luciferase activity that was similar in speed when comparing HdV^wt^-Nluc to HdV^ko^-Nluc, as was expected. In contrast, the ActD-imposed block to transcription resulted in a much faster drop in luciferase activity for HdV^wt^-Nluc compared to HdV^ko^-Nluc transfected cells, with the data from the first 4 hours suggesting an effective translational half-life of 64 and 445 minutes respectively. This more rapid drop for HdV^wt^-Nluc seen after ActD-treatment confirmed that ribozyme activity was destabilizing translationally active mRNA.

To directly examine mRNA-embedded HdV ribozyme activity and establish whether it might be influenced by translational activity, cells were transfected with the HdV^wt^-Nluc or HdV^ko^-Nluc vector, and treated with either ActD alone, or ActD plus CHX. Transcript integrity and abundance was then monitored over a period of 4 hours using Northern blot (Fig. 3d). Interestingly, in addition to the ~1.4 kb full length transcript seen in all transfected cells, a putative 1.1 kb HdV ribozyme cleavage product was also observed in the HdV^wt^-Nluc transfected cells. The appearance of this extra RNA species, coupled to the fact that the HdV^wt^-Nluc full length transcript disappeared over time whereas the levels of HdV^ko^-Nluc stayed more or less unchanged, confirmed that the HdV^wt^ ribozyme was cleaving and destabilizing the full length transcript. Importantly, the rate at which the HdV^wt^-Nluc transcript disappeared after ActD treatment was the same irrespective of whether CHX was present or absent, indicating that translational activity did not significantly influence ribozyme cleavage.

### Validating the ability of the reporter system to detect differences in HRV protease mediated shut down of gene expression

To confirm that the reporter system was able to detect differences in the rate of shut down of gene expression by HRV proteases, we made use of a bidirectional promoter contained within the A16^TAG^(+/+) and (−/−) vectors to generate constructs expressing both the HRV proteases and HdV^wt^-Nluc. Cells were electroporated with either 2, 5 or 10 μg of these two constructs, balancing the total amount of DNA introduced into cells through use of the A16^TAG^(−/−) plasmid lacking any Nluc cassette. Luciferase activity was then monitored hourly. At the 1 hour time point luciferase values positively correlated with amount of HdV^wt^-Nluc vector electroporated into cells (Fig. 4a). Importantly, there was no obvious difference in the signals between the A16^TAG^(+/+) and (−/−) experimental groups, indicating that the proteases had yet to exert an impact on gene expression. For this reason, all subsequent time point values were normalized to their 1 hour counterparts to correct for transfection efficiency and to allow for more effective comparison to be made. Interestingly, when looking at these later time points, a clear difference was seen in luciferase values when comparing between the different (+/+) experimental groups (Fig. 4b). Consistent with having reduced protease expression from transfecting lower amounts of plasmid, reducing the amount of A16^TAG^(+/+) vector transfected into cells resulted in higher luciferase peak values and a delay in reaching these peak values. These differences in luciferase values reached significance by 2 hours when comparing across the three groups and remained significant for the remainder of the assay. In contrast to the A16^TAG^(+/+) results, the A16^TAG^(−/−) experimental groups showed a more prolonged increase in luciferase activity over time that reached higher peak levels at later time points (Fig. 4c). More importantly, the amount of vector used made no significant difference to the normalized luciferase values at any of the time points over which the 5 hour experiment was run. This confirms that the differences seen between the different A16^TAG^(+/+) experimental groups are exclusively due to varying protease activity within transfected cells.

**Figure 4.**
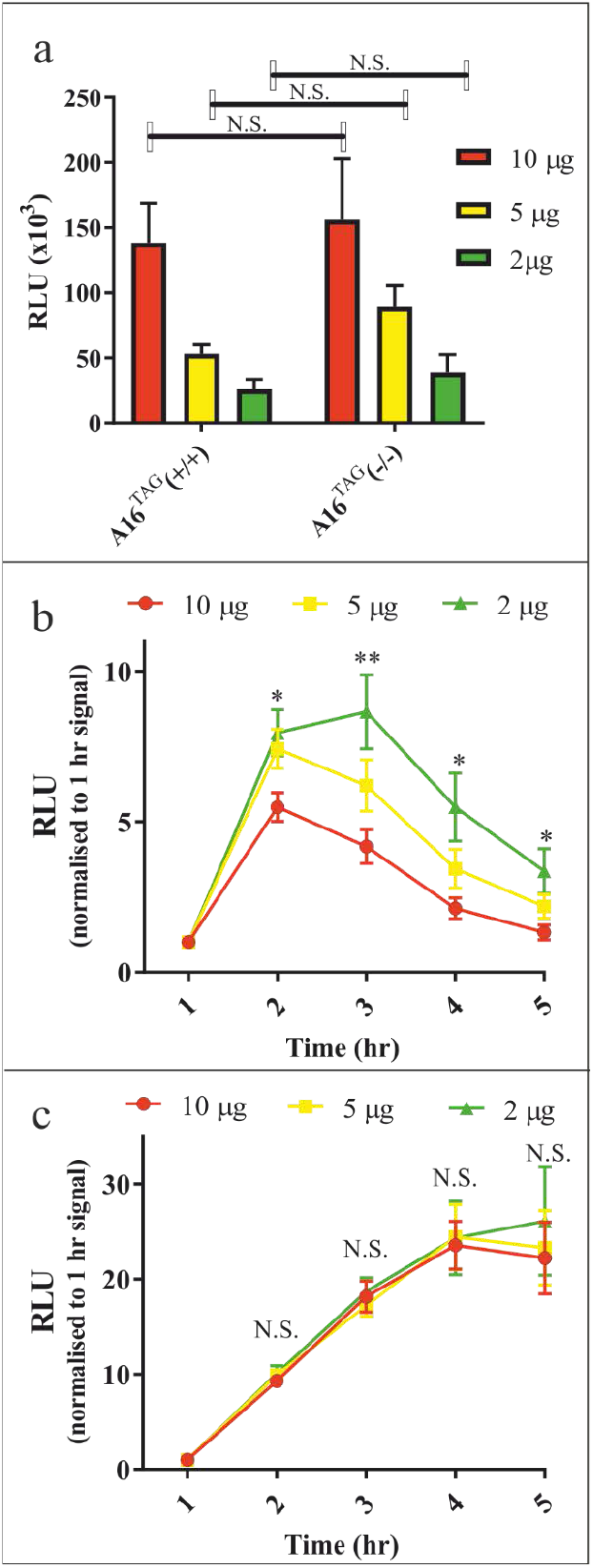
Transfecting different quantities of a vector expressing the active form of the HRV proteases impacts on the rate at which Nluc activity is inhibited. Cells were electroporated with 2, 5 or 10 μg of a plasmid vector expressing A16 2A and 3C epitope tagged proteases ((+/+) or (−/−) versions) and the HdV^WT^Nluc reporter. The total amount of DNA electroporated in each experimental group was adjusted to 10μg by the addition of an A16^TAG^(−/−) vector that lacked the HdV^WT^Nluc coding region. Graphical representations include (a) luciferase values 1 hour post transfection, as well as all subsequent time point values for (b) A16^TAG^(+/+) and (c) A16^TAG^(−/−) after normalizing to the 1 hour transfection values. Data shown represents the mean ± S.E.M. of 4 separate experiments. The existence of a statistical significance difference when comparing across the 3 experimental DNA concentrations are indicated (* = p<0.05, ** = p<0.01, N.S. = not significant).

### Comparing the ability of A16, B4 and C2 proteases to shutdown gene expression

Having validated the HdV^wt^-Nluc reporter system, the cassette was cloned into the B4^TAG^(+/+) and C2^TAG^(+/+) containing vectors and these along with their A16^TAG^(+/+) and A16^TAG^(−/−) counterparts were electroporated into cells. Monitoring luciferase values over time revealed that there was no difference in signals at 1 hour post transfection (Fig. 5a), again allowing the data to be normalize to this time point. At 2 hours post transfection, no significant difference was observed among the 4 experimental groups, but from 3 hours onwards luciferase values from the protease inactive A16^TAG^(−/−) construct continued to rise and were significantly higher than the three other protease active constructs which instead had declining luciferase values (Fig. 5b). When comparing between the different protease active constructs there was also a trend at 3 and 4 hours for A16^TAG^(+/+) to suppress luciferase activity more than B4^TAG^(+/+) and C2^TAG^(+/+), although this did not reach significance.

**Figure 5.**
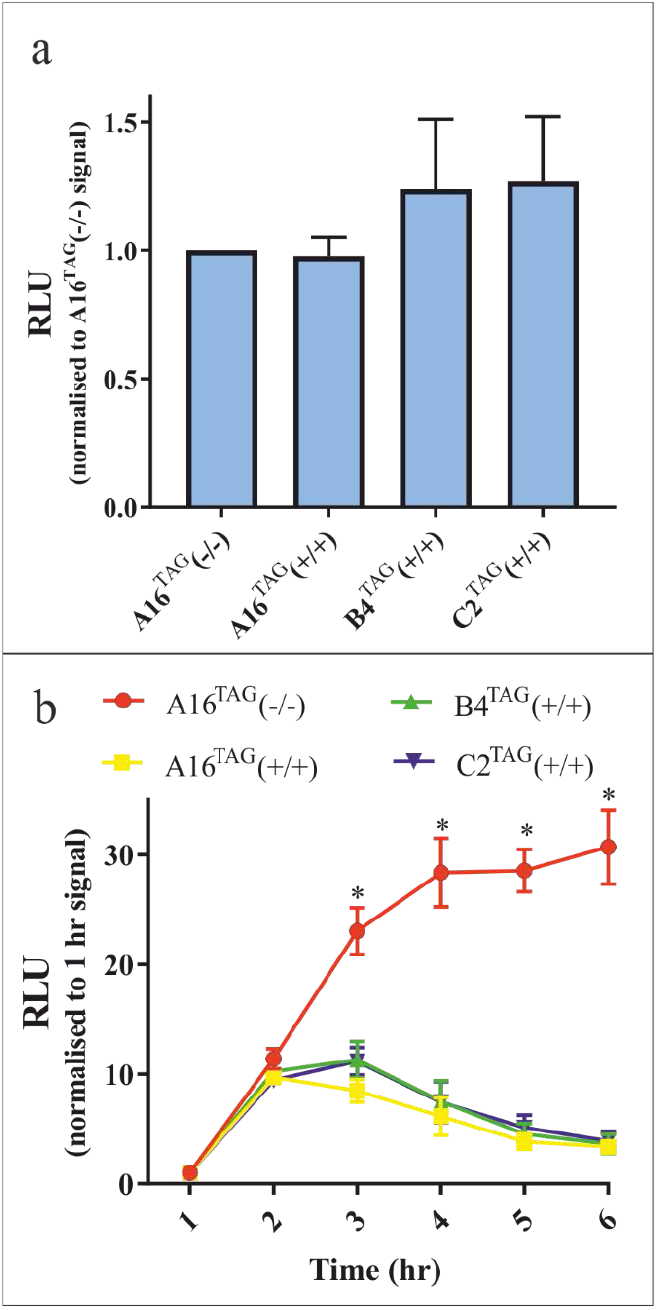
Comparison of the different tagged active protease vectors to block gene expression. Cells were electroporated with vectors co-expressing both HRV proteases and HdV^WT^Nluc, and luciferase activity monitored over time. Graphs shown represent (a) data for luciferase values 1 hour post transfection, as well as (b) time point values for the different experimental groups after normalizing to the 1 hour transfection values. Data shown represents the mean ± S.E.M. of 5 separate experiments. Statistical significance between A16^TAG^(−/−) and all other groups (p<0.05) is indicated by an asterisk. No statistical difference existed between the different experimental groups 1 hour post-transfection.

Although N-terminal epitope tagging of HRV proteases is a strategy that has been used in previous cell based studies[10, 32] and we had shown that our tagged proteases were active, it was nonetheless important to establish what were the consequences when the proteases were expressed in their native form. We therefore modified the existing protease expression vectors such that the first codon of 2A and 3C was placed immediately downstream from the ubiquitin coding regions they were linked to, generating A16^NT^(+/+), B4^NT^ (+/+) and C2 ^NT^(+/+) constructs. Western blot analysis of cells transfected with these constructs (Fig. 6a) confirmed that that GFP was present and that both eIF4GI and AUF1 cleavage occurred, demonstrating that both 2A and 3C were being produced in an active form. Transfection of the equivalent constructs expressing the HdV^WT^Nluc reporter was then used to assess the rate at which new gene expression was inhibited by these tag-free proteases. Similar to previous experiments, the luciferase signal at 1 hour was the same when comparing across the 4 experimental groups (Fig. 6b), allowing data to be normalized to this first time point reading. Interestingly, in contrast to the results from the protease active tagged constructs, a difference was seen between A16^TAG^(−/−) and all untagged protease active constructs at 2 hours which reached significance for A16^NT^(+/+) and C2 ^NT^(+/+) (p<0.05) and was almost significant for B4^NT^ (+/+) (p=0.053) (Fig. 6c). For all subsequent time points the luciferase activity from all active protease constructs was significantly different from that produced by the A16^TAG^(−/−) control. More importantly at 2 hours a trend a started to emerge when comparing between the three (+/+) constructs similar to that seen for the tagged (+/+) constructs, with A16^NT^(+/+) suppressing luciferase activity more effectively than B4^NT^(+/+) and C2^NT^(+/+) (Fig. 6c). Importantly, this difference reached significance at the 3 hour time point before disappearing at later time points (Fig. 6d), suggesting the A16 proteases were more rapidly shutting down early gene expression compared the B4 and C2 proteases.

**Figure 6.**
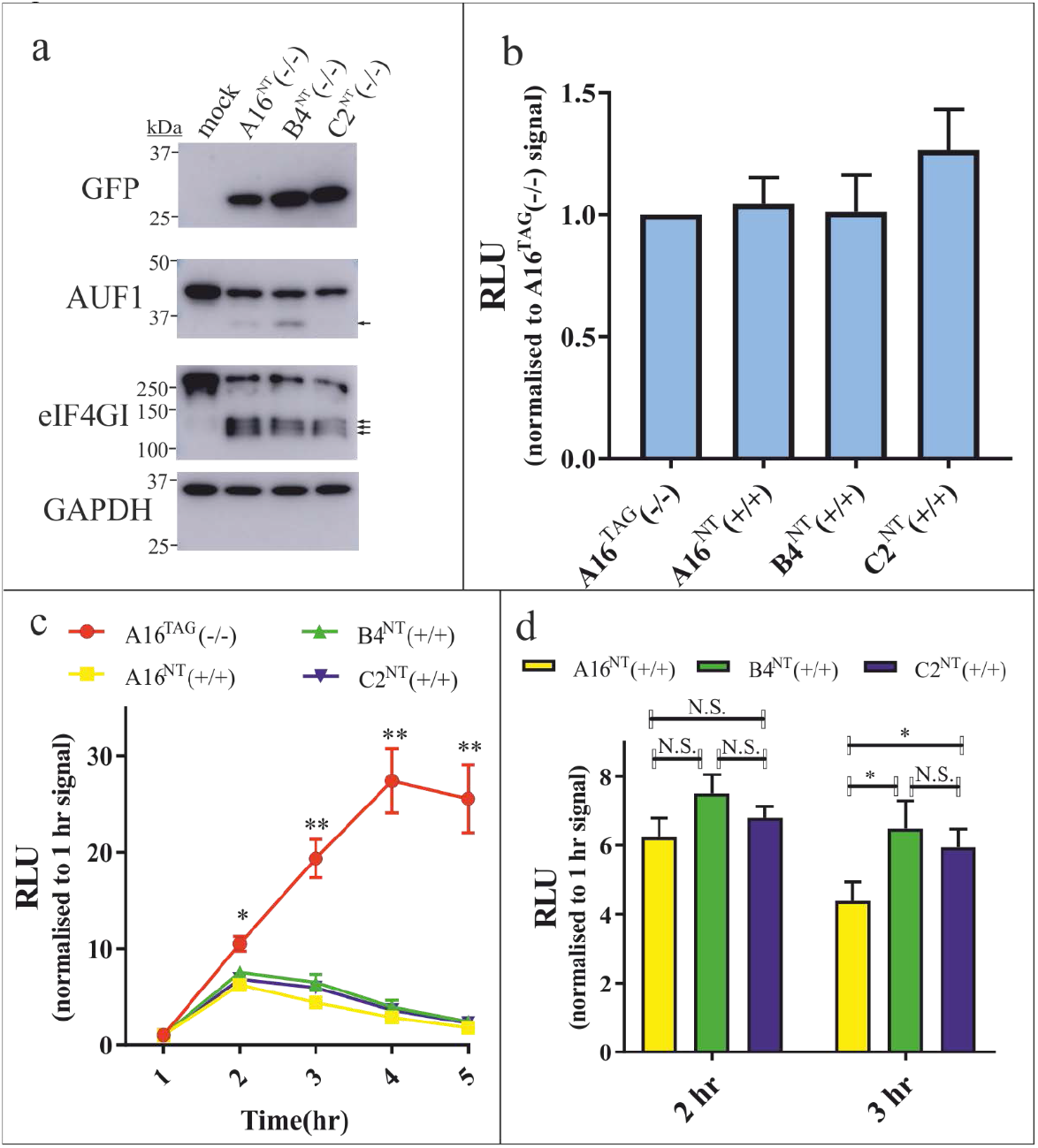
Expressing non-tagged HRV proteases in cells and their impact on gene expression. (a) Constructs expressing 2A and 3C lacking an N-terminal epitope-tagged extensions were transfected into cells which were subsequently analysed by Western blot. Arrows indicate protease cleavage products. (b-d) Comparable constructs to those in (a) but co-expressing HdV^WT^Nluc were electroporated into cells and analysed for luciferase expression. Graphs show (b) data for luciferase values 1 hour post transfection, as well as (c) all time point values for the different experimental groups after normalizing to the 1 hour transfection values. A subsection of this latter data (d) is presented as a bar graph to illustrate differences between the protease active constructs at 2 and 3 hours. Time points where pairwise comparisons show statistically significant differences (p<0.05) between A16^TAG^(−/−) and both A16^NT^(+/+) and C2^NT^(+/+) (*), or A16^TAG^(−/−) and all three protease active constructs (**), are highlighted in the graph shown in (c). Statistically significant differences between the protease active constructs (* = p<0.05) are highlighted in the graph in (d). Data represents the mean ± S.E.M. of 7 separate experiments. No statistical difference existed between the different experimental groups 1 hour post-transfection.

### Comparing the ability of 2A and 3C to shut-down early gene expression

To determine the extent to which 2A and 3C contributed to the early shutdown of gene expression, modified versions of the A16^TAG^(+/−) and (−/+) constructs carrying the HdV^WT^Nluc cassette were electroporated into cells and luciferase activity monitored over a 7 hour window (Fig. S4). Remarkably, A16^TAG^(+/−) inhibited gene expression as effectively as A16^TAG^(+/+), there being no apparent difference between these two experimental groups for any of the time points examined. In contrast, inhibition of luciferase expression by A16^TAG^(−/+) compared to the A16^TAG^(−/−) control was negligible for up to 5 hours post transfection, and even by 7 hours still showed only a marginal reduction in activity.

3C is normally targeted to the nucleus at early time points during infection as a result of being expressed as a 3CD precursor, with a nuclear localization sequence (NLS) in 3D facilitating this process. While 3C’s size was expected to enable the protease to access the nucleus as a result of passive diffusion through the nuclear pore, we none-the-less wondered whether failing to actively target it to the nucleus meant its ability to shut down gene expression was being underestimated. Based on our earlier observations that a A16^NT^(+/+) was more effective at shutting down gene expression compared to A16^TAG^(+/+), we also wanted to ensure that the amino-terminal epitope tags were not confounding our assessment of 3C’s activity. Therefore, the HdV^WT^Nluc reporter version of the A16^NT^(+/+) construct was modified to express epitope tag-free versions of an inactive 2A with an active 3C (A16^NT^(−/+)), an inactive 2A with an active 3C containing a C-terminal NLS (A16^NT^(−/+^NLS^)), an active 2A with an inactive 3C (A16^NT^(+/−)) and an active 2A with an active 3C containing a C-terminal NLS (A16^NT^(+/+^NLS^)). These constructs were transfected into cells along with A16^NT^(+/+) and A16^TAG^(−/−) controls and luciferase monitored over a 7 hour period (Fig. 7a and b). Consistent with earlier experiments using tag-free constructs, while luciferase levels were broadly comparable across the different experimental groups 1 hour post transfection (Fig. 7a), this changed after 2 hours. Importantly, at this time point it was only the cells transfected with constructs expressing an active 2A where luciferase levels were seen to drop compared to cells transfected with the A16^TAG^(−/−) control (Fig. 7b). Equally noticeable was the fact that when 2A was active, the presence or absence of an active 3C made no appreciable difference to luciferase levels at any of the time points, irrespective of whether 3C possessed an NLS extension or not. Indeed, when 2A was inactive but 3C was active, a drop in luciferase activity compared to the A16^TAG^(−/−) control only started to become appreciable towards the end of the time period, and the presence or absence of a NLS on 3C made little or no difference to this trend. To confirm that the NLS extension on 3C was targeting it to the nucleus, a A16^TAG^(−/−^NLS^) construct expressing epitope tagged inactive forms of both proteases but where 3C also possessed the NLS tag, was transfected into cells. Immunofluorescence analysis of these cells alongside cells transfected with A16^TAG^(−/−) where 3C lacked the NLS extension, verified that an NLS-containing 3C was more effectively targeted to the nucleus (Fig. 8). Based on this accumulated data using both epitope tagged and non-tagged constructs, we conclude that 2A plays an almost exclusive role in shutting down new gene expression.

**Figure 7.**
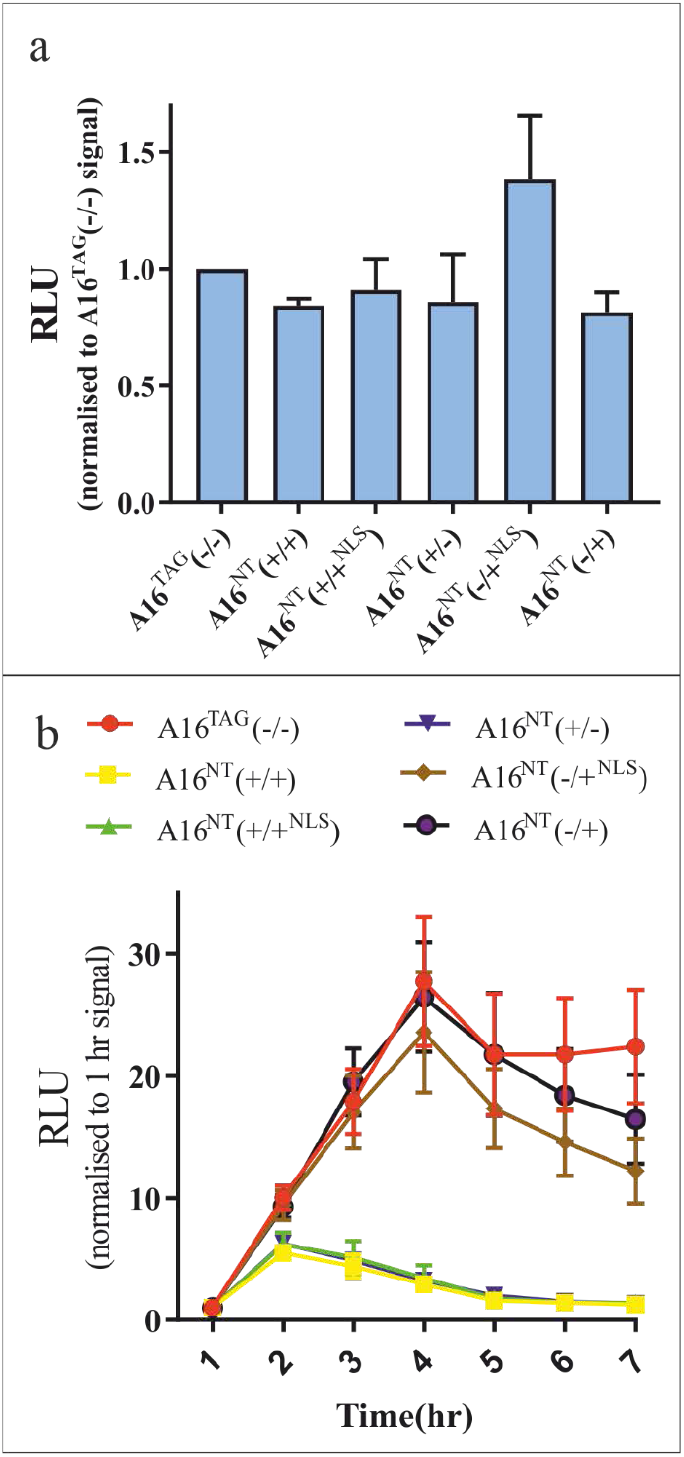
Assessing the individual contribution made by 2A and 3C on the early shutdown of gene expression. Cells were electroporated with vectors expressing combinations of active and inactive non-tagged A16 2A and 3C proteases as well as the HdV^WT^Nluc reporter. A comparable A16^TAG^(−/−) construct was included as a control. (a) Data for luciferase values 1 hour post transfection, and (b) time point values for the different experimental groups after normalizing to the 1 hour transfection values. Data shown represents the mean ± S.E.M. of 4 separate experiments.

**Figure 8.**
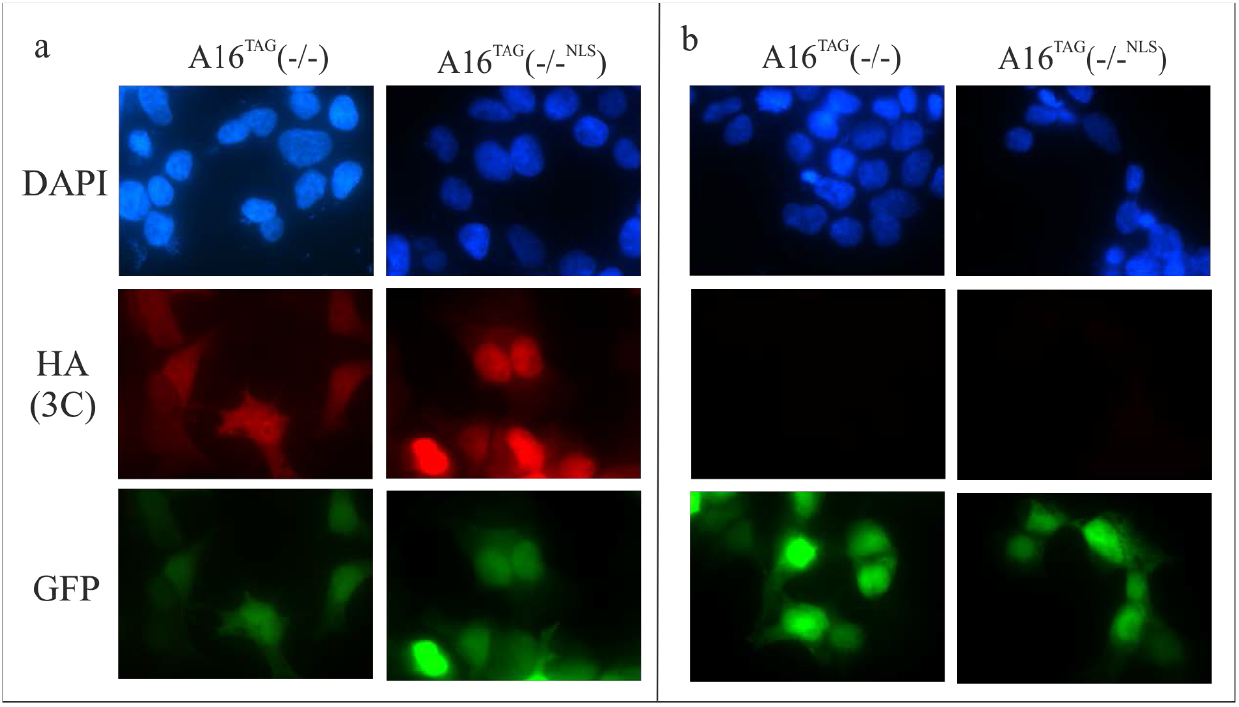
Visualizing the location of 3C by immunofluorescence microscopy when expressed with and without an NLS tag. Cells were transfected with constructs expressing epitope-tagged versions of inactive 2A and 3C proteases from A16 and subsequently fixed and stained with (a) a mouse anti-HA mAb followed by a TRITC-conjugated anti-mouse secondary antibody or (b) a TRITC-conjugated antimouse secondary antibody only.

## Discussion

To our knowledge, this is the first study to examine whether the combined activity of 2A and 3C from different HRV species differ in capacity to block new gene expression. It is also the first to assess the relative contribution each protease makes to achieving this goal. While analysis was restricted to a single isolate from each HRV species, we nonetheless found that there were differences between isolate proteases. It is almost certain that there are many viral factors, in addition to protease-directed inhibition of gene expression, that contribute to HRV pathogenicity, including receptor usage[33–37] and viral replicative capacity at both upper respiratory (34°C) and lower respiratory (37°C) tract temperatures[38]. Nonetheless, given our observation that HRV-A proteases appear more effective at inhibiting new gene expression than the proteases from other HRV species, it is worth noting that it is this same species, along with HRV-C, that have been suggested by some studies to be associated with severest disease[39–41].

Interestingly, it took substantially longer to observe an effect on gene expression in the presence of active 3C versus active 2A. After plasmid transfection, the latter protease started to exert an effect on reporter gene expression with 3 hours, and by ~5 hours inhibition reached near maximal levels (~85% inhibition compared to protease inactive control plasmid). In contrast, 3C failed to have a noticeable impact on reporter gene expression until 6 hours, and even at this point inhibition was marginal (Fig. 6b). Indeed, 3C seemed to have a minimal impact above and beyond that imposed by 2A, as we also failed to see any noticeable difference in inhibition of reporter gene expression when only 2A activity was present in the cell compared to when the activity of both proteases were present. Of course, our data do not provide an accurate estimate of the window of time between when the two proteases impact on gene expression in an infected cell, given that the rate of protease expression in our pol II-based assay will be different to that of the former scenario. For instance in an infected cell, once 2A inhibition of cap-dependent translation is complete, IRES-dependent translation would increase and drive more rapid production of both 2A and 3C[42]. Infected cells would also experience a rapid and continuous increase in viral transcript which would likely drive heightened levels of protease expression beyond that achievable using transfection. Both these two factors could well shorten the window of time between when 2A inhibits gene expression and when 3C starts to have an effect. Nonetheless, whatever timescales operate during HRV infection, 2A would appear to inhibit host gene expression considerably earlier than 3C. This observation both parallels and complements a recent study, published during the preparation of this manuscript, showing that 2A rather than 3C is responsible for blocking the intrinsic antiviral response in enterovirus-infected cells[43]. What then is the selective advantage gained by 3C targeting processes involved in gene expression, and most specifically transcription, given the evidence that this activity is important for viral replication[8]? Several possibilities present themselves. Firstly, while HRV 3C cleaves Oct-1[21], most of the studies looking at the effect of enterovirus infection on mRNA transcription and its importance have been done with poliovirus. It is therefore possible that HRV does not actively target mRNA transcription in the same way as poliovirus. Consistent with the view point that HRV might not depend on rapid shut-down of mRNA transcription is the recent observation that unlike poliovirus, HRV fails to use 3D to suppress mRNA splicing[44]. A second possible explanation relates to the observation that while most gene transcription is quickly suppressed by enteroviruses, a minority of transcripts – some with antiviral activity – are both upregulated and translated[45, 46]. If one assumes 2A is responsible for the more widespread global inhibition of transcription and translation, 3C may be acting to help suppress those genes that bypass 2A inhibition. Alternatively, inhibition of transcription by enteroviruses may not be principally aimed at blocking protein expression, but instead may serve to prevent production of RNA transcripts with direct antiviral functions such as microRNAs[47]. Blocking nuclear RNA transcript production more generally might also enhance ribonucleotide availability for viral genome replication. Finally, it is possible that 3C inhibition of gene transcription is more restricted when expressed as a mature product rather than a 3CD precursor. Selective subcellular targeting alone seems unlikely to be able to facilitate such a restriction, as we found that active targeting of 3C to the nucleus made little difference on its ability to shut-down gene expression. However, alternative possibilities are that 3D binds host proteins thus facilitating interaction with 3C, or 3D subtly alters 3C substrate recognition. There is certainly precedence for such ideas, given that 3C cleavage of the enterovirus capsid protein[48] and the TIR-domain-containing adapter-inducing interferon-β [49] is dependent on enterovirus and hepatitis A virus 3C being expressed as a 3CD precursor. [44]. It would be interesting to know whether transcription was inhibited more rapidly as a result of expressing 3C or 3CD, assuming it was possible to achieve balance expression levels and target these proteins to the nucleus.

A technical development that made this study possible was the generation of a reporter construct that produced a highly destabilized mRNA, allowing rapid detection of changes to gene transcription, mRNA maturation and mRNA export. While the use of cis-acting ribozymes to control gene expression is a strategy that has been used by others [50, 51], to our knowledge this is the first study to use a ribozyme as substitute to more traditional mRNA destabilizing approaches for the purposes of producing a reporter mRNA with a shortened translational half-life. Selection of the HdV ribozyme in this study over the more often favoured hammerhead ribozyme was due to the former RNA exhibiting a slow rate of folding[52]. Both this feature, combined with the introduction of an intron within the HdV ribozyme so as to deposit an exon-junction complex on the ribozyme sequence, were expected to minimize mRNA cleavage prior to nuclear export. A clear advantage to using an HdV ribozyme in this way is that its impact on mRNA stability is not subject to regulation, unlike that of endogenous RNA destabilizing pathways such as recognition of AU-rich elements and nonsense-mediated decay. Importantly both these two decay pathways are modulated/usurped in virus-infected cells, potentially compromising their ability to be used in studies such as ours [30, 31, 53–56]. Given that the apparent half-life of our reporter signal after transcription termination compares favourably with that of other dynamic reporter systems[57], we believe it offers certain benefits that may make it useful for other studies.

Epitope tagging of proteins is an effective means of monitoring expression levels in cells when other suitable immunological reagents are lacking, and has been used in the past for studies looking the impact of 2A and 3C in cells [10, 32]. We found that while tagged proteases continued to display activity towards their respective substrates, 2A had diminished activity regards the early inhibition of reporter gene expression. Processing of the N-terminus of 2A is a cis-acting event, demanding that carboxy-terminal end of VP1 lies across the active site of 2A prior to cleavage. We therefore suspect that the epitope tag is sterically hindering substrate access to the active site of 2A in a way that uniformly suppresses protease activity for all 3 species. Consistent with this notion, the trend in inhibition of host gene expression by the three different HRV species proteases remained the same, irrespective of whether the proteases were tagged or not, with HRV-A16 being more active than B4 or C2. Future studies need to be aware of the potential problems using amino-terminal epitope tagged 2A constructs when assessing protease activity.

In conclusion, natural variation in rhinovirus protease activity translates into measurable differences in the speed at which the proteases are able to block gene expression from the infected host cell nucleus. This almost certainly will influence pathogenicity, although the extent to which it does so remains to be determined. Further work is needed to establish whether the activity of proteases from a larger sample of HRV isolates follow the same pattern as observed in this study, and determine whether such variation relates to clinical outcome.

## Abbreviations

ActD: actinomycin D
AUF1: AU-rich element RNA-binding protein 1
CHX: cycloheximide
DAPI: 4,6-diamidino-2-phenylindole
EMCV: encephalomyocarditis virus
HCV: hepatitis C virus
GAPDH: glyceraldehyde 3-phosphate dehydrogenase
HA: hemagglutinin
HdV: hepatitis delta virus
HRV: human rhinovirus
IRES: internal ribosome entry site
Nluc: nanoluciferase
NLS: nuclear localization signal
Nup: nucleoporin
RLU: relative light units
VSV: vesicular stomatitis virus

## Acknowledgements

We are grateful to Prof Ken Hirasawa (Memorial University, Canada) for the gift of pR.EMCV.F.

## Author’s contributions

DS: investigation, data curation, writing-review and editing. IF: investigation, resources. BS: resources. DD: conceptualisation, writing-review and editing, project administration, funding. CM: conceptualisation, methodology, data curation, investigation, formal analysis, writing – original draft preparation, visualisation, supervision, project administration, funding.

---

**Figure S1.**
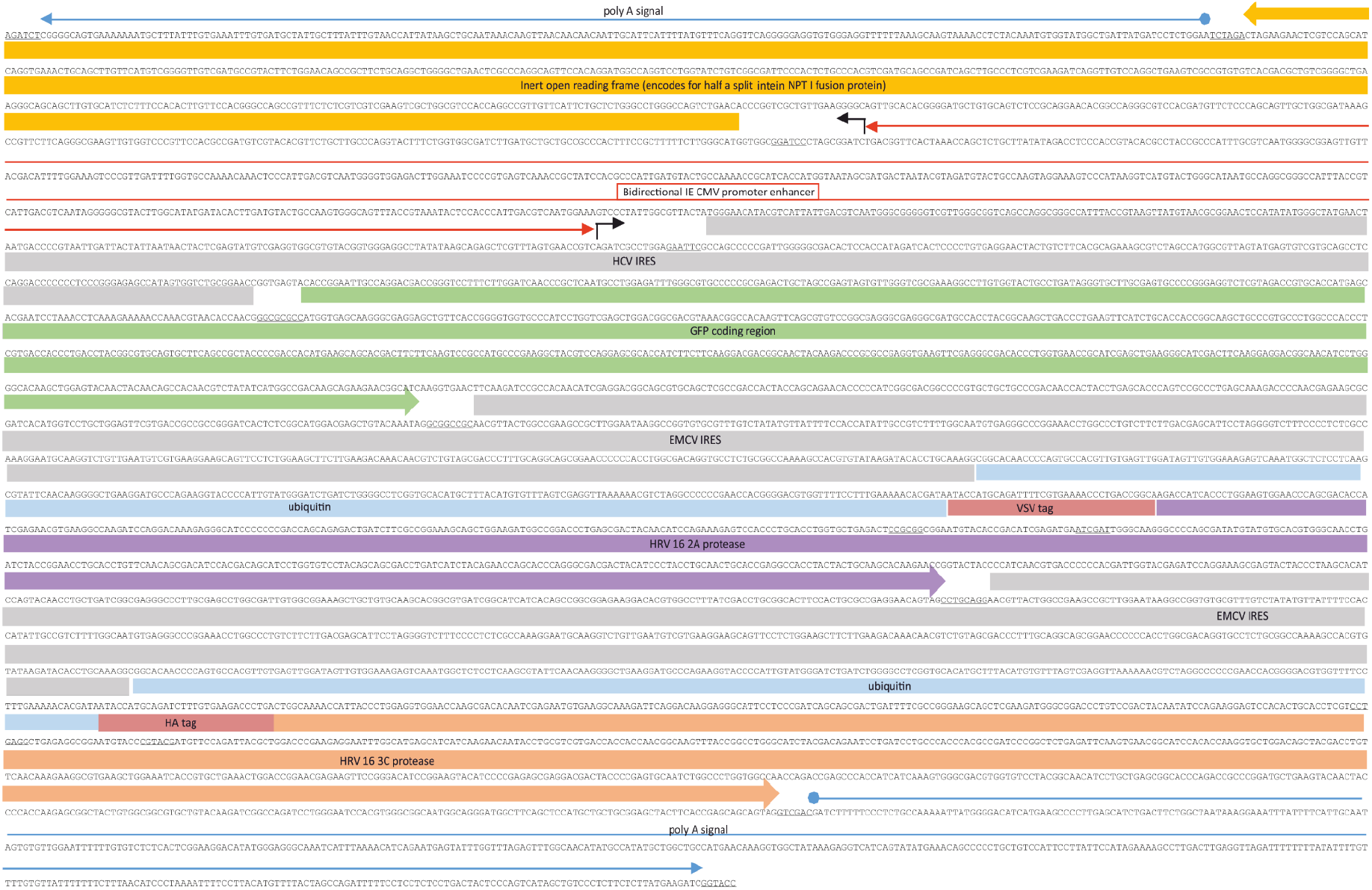
Sequence of the dual promoter mammalian expression cassette used for expression of A16^TAG^(+/+) mRNA. Sequences of restriction sites used for cloning during the construction of this and subsequent pCIPEP vectors are underlined, with the cassette itself cloned into LITMUS28 via BglII and KpnI sites found at its extreme 5’ and 3’ ends.

**Figure S2.**
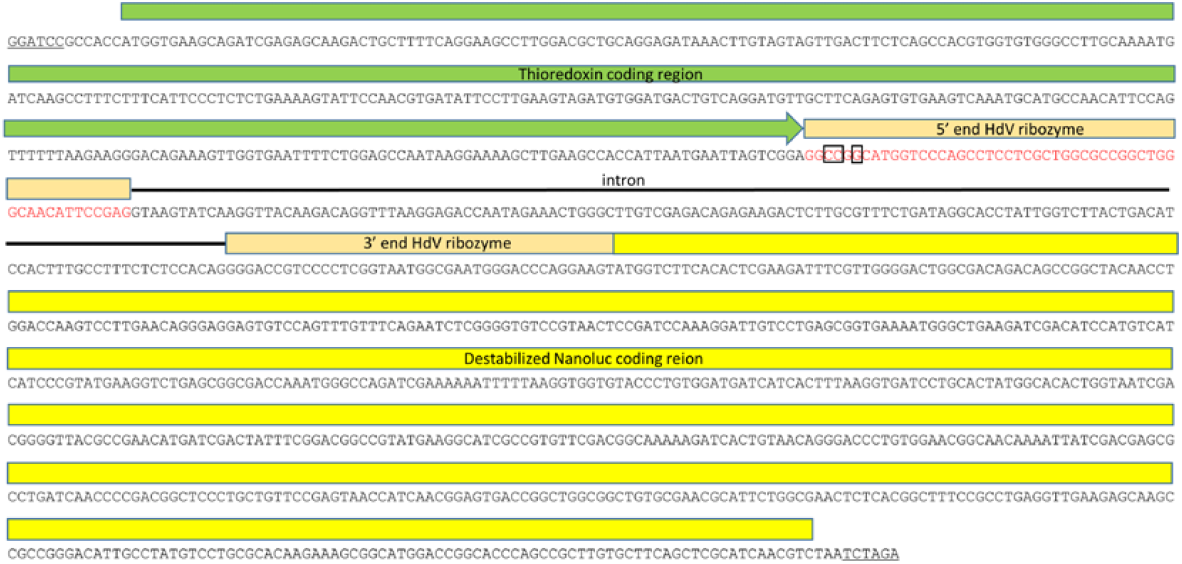
Sequence of HdV^WT^Nluc, the Nluc reporter cassette carrying an intron split-HdV ribozyme. Boxed nucleotides are those targeted for synonymous mutagenesis and changed to adenine bases to generate HdV^KO^Nluc, an equivalent reporter cassette carrying an inactive ribozyme.

**Figure S3.**
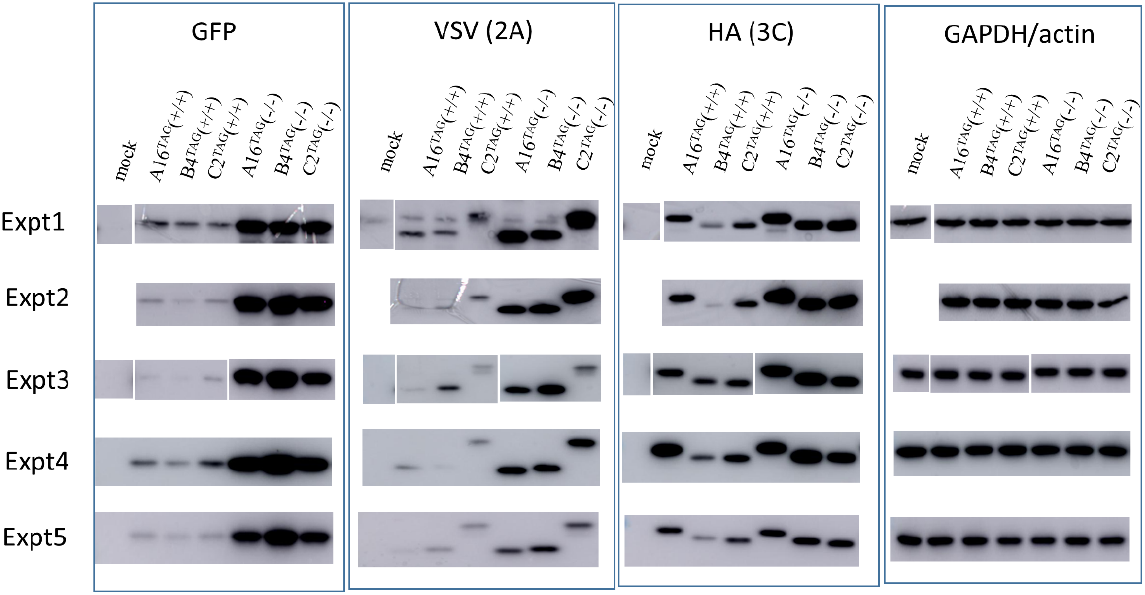
Western blot of cells transfected with the epitope-tagged protease expression vectors encoding the 2A and 3C proteases from A16, B4 and C2. The results from 5 independent transfection experiments are shown. Blots from experiment 1 and 3 have been cropped and edited to re-order the lanes so that they match those of the other blots. Blots from experiment 2 did not include a mock control lane.

**Figure S4.**
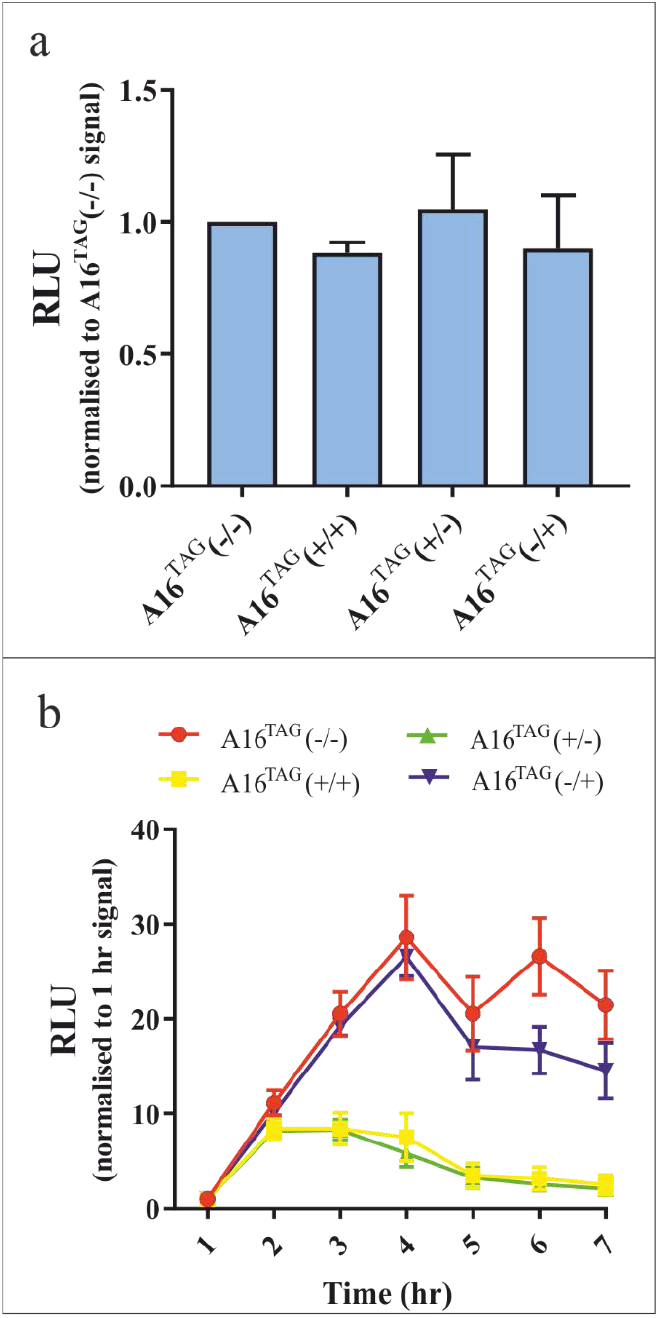
Assessing the individual contribution made by epitope tagged 2A and 3C on the early shutdown of gene expression. Cells were electroporated with vectors co-expressing combinations of active and inactive A16 epitope tagged 2A and 3C proteases as well as the HdV^WT^Nluc reporter, and luciferase activity monitored over time. (a) Shows the data for luciferase values 1 hour post transfection. (b) Shows time point values for the different experimental groups after normalizing to the 1 hour transfection values. Data shown represents the mean ± S.E.M. of 3 separate experiments.

